# Simple, fast, low-cost method to produce transgenic pollen

**DOI:** 10.1101/2025.05.26.655774

**Authors:** Hui Wang, Yujie Zhang, Wen Qu, Xin Zhang, Haiyong Qu

## Abstract

There are two main technical bottlenecks in plant genetic transformation: the intracellular delivery of exogenous genes and tissue culture. This study is an elaboration of a method combining needle-free jet injection (NFJI) and vacuum infiltration technology to efficiently introduce exogenous genes into pear (Rosaceae) plant pollen. This method was verified with the model plant tobacco (Solanaceae) wherein the GFP reporter gene was successfully introduced into the pollen and expressed. Then, the gene was transferred to the seeds via pollination, and after the seeds were sown, gene expression was observed at the roots of the seedlings. Furthermore, in this study, the apple (Rosaceae) gene MdGH3.1 was introduced into tobacco using NFJI, resulting in transgenic seedlings that were shorter than the seedlings in the control group and presented fewer, shorter root hairs, confirming that the MdGH3.1 gene was functional. In addition, the applicability of this method to other plant species, including dicotyledonous and monocotyledonous plants, was explored. In particular, when the MdGH3.1 gene was introduced into Arabidopsis (Brassicaceae), the phenotype of the resulting plants was consistent with that of the transgenic tobacco. The results indicate that genes can be effectively delivered into plants by this method without on the need for tissue culture. Moreover, the instruments used in this method are portable, affordable, and easy to operate, allowing for direct utilization in the field, indicating the great application potential of this method.

## Introduction

Agriculture has reached an important technological turning point in its development and is facing severe, unprecedented challenges caused by population growth and global climate change (Chen et al., 2022). The speed and scope of plant breeding programs are set to undergo fundamental changes due to significant breakthroughs in genetic modification technology and genome editing and the rapid emergence of new strategies for bioengineered crop breeding, which thereby increase our ability to ensure an adequate food supply and support population growth (Li et al., 2024). Despite recent advances in genetic modification technology and CRISPR editing, efficiently delivering gene modification components into plant cells in the first step of plant transformation and genome editing, remains the main bottleneck that limits the widespread application of these technologies (Lu et al., 2024; Wang and Doudna, 2023). Therefore, developing innovative methods to ensure high gene transmission efficiency is particularly important (Pacesa et al., 2024; van Haasteren et al., 2020). Since the 1980s, two main methods of gene transfer have been used in higher plants: direct gene transfer and Agrobacterium-mediated transformation (Chen et al., 2022). Direct gene transfer methods are considered effective strategies to overcome the potential barriers in transformation competence. Traditionally, particle bombardment, also known as biolistics, has been used to facilitate gene delivery, but this process requires expensive equipment (Ozyigit and Yucebilgili Kurtoglu, 2020). Agrobacterium-mediated transformation gene delivery systems are expensive but display considerable transformation efficiency and can transfer large DNA fragments to plant chromosomes; therefore, this method is preferred for plant transformation (Mei et al., 2024). However, only a few plant host genotypes can accept Agrobacterium infection and the Agrobacterium infection efficiency is low in most plants, which limits its widespread application to some extent (Khatun et al., 2022; Rahman et al., 2024). The regenerative capacity of plants (i.e., the ability to form fertile plants from transformed cells) is another key barrier to obtaining transgenic plants. Traditional methods mainly induce callus formation from tissue explants to produce transgenic plants. However, callus formation is a time-consuming and labor-intensive process (Chen et al., 2022). Germ cell transformation does not rely on regeneration and mainly targets female (egg cells) or male (sperm) gametes. Transformation is carried out before gamete fusion and the production of diploid zygotes (Anjanappa and Gruissem, 2021). In general, technologies that do not rely on regeneration are often considered more efficient than those that do (Bélanger et al., 2024). Pollen grains have significant advantages as carriers for introducing exogenous genes into a germ line, as they are easily isolated and abundant and their transformation is convenient. However, the pollen grain surface is covered with an exine layer derived from the anther tapetum and a cell wall composed of a thick outer layer and a thin inner layer, and their presence limits the transfer of exogenous genes into pollen (Levengood et al., 2024). To solve this problem, researchers have employed various techniques to genetically modify pollen; however, these techniques require fresh pollen or even anthers (Wang et al., 2022; Zhao et al., 2017). For woody plants for which stable genetic transformation systems are difficult to establish, such as pear, apple, and peach in the Rosaceae family, the flowering period usually lasts only approximately 7 d. During artificial pollination, pollen from the previous year, which has been preserved under low-temperature and dry conditions, is often used. Thus, using fresh pollen is a particularly prominent challenge (Yin et al., 2021).

Therefore, despite the developments in genetic modification technologies, transformation protocols should be optimized with a particular focus on simplifying procedures, increasing operational convenience, and facilitating breeding (Cunningham et al., 2018). Needle-free jet injection (NFJI) devices are portable and use a high pressure source to expel liquids at speeds exceeding 100 m/s for epidermis and dermis penetration, thereby effectively delivering the material to the subcutaneous or muscle tissues (Mao et al., 2023; Ravi et al., 2015; Schoppink et al., 2024). During Agrobacterium-mediated transformation, a vacuum is applied to create negative pressure, which leads to a reduction in the volume of air outside the cell membrane, thereby promoting the efficiency of Agrobacterium invasion of host cells (Mithra et al., 2017; Shi et al., 2024). In this study, NFJI was used to deliver plasmids containing exogenous genes into pear (*Pyrus* spp.) pollen. Then, the pollen was placed under vacuum to facilitate plasmid entry into the pollen, after which the exogenous genes were expressed in the pollen. The effectiveness of this technique was validated using tobacco (*Nicotiana benthamiana*) as a model plant. Currently, there have been no reports of using NFJI to deliver genetically modified components.

## Results

This study utilized NFJI (Supplementary figure S1) to spray ‘Akitsuki’ pollen with distilled water and then employed scanning electron microscopy to observe the pollen grain surfaces. The density of the pores on the sprayed pollen surface was significantly greater than that of the untreated control pollen (Figure 1A-C). Notably, this treatment did not inhibit but rather promoted both the pollen germination rate and pollen tube elongation (Supplementary figure S2). Using NFJI and an easy to carry vacuum pump (Supplementary figure S3), the GFP gene was delivered into the pear pollen following the process shown in Figure 1D, after which the pollen was cultured for 2 h. No fluorescence signals were observed in the control group or the pollen tubes treated via vacuum infiltration (Figure 2A and B); however, the pollen grains exhibited fluorescence, a phenomenon attributed to the autofluorescent properties of the pollen grains themselves (Dunker et al., 2021). Huang et al. (2014) reported that GFP can be used as a reporter gene to effectively detect if a gene was successfully introduced into pollen cells. These authors suggested that GFP gene transfer can be considered successful only when both the pollen grains and pollen tubes fluorescence (Huang et al., 2014). The results of this study indicate that exogenous genes cannot be effectively introduced into pear pollen cells via vacuum infiltration alone. However, after performing the treatment process shown in Figure 1D, fluorescence signals were observed in the pollen tubes (Figure 2C). Additionally, we altered the process shown in Figure 1D by adopting two other methods (Figure 2D and E). The pollen tubes processed by these two altered methods also exhibited fluorescence, but the gene transformation rates and fluorescence intensities of the pollen tubes were both lower than those observed with the method presented in Figure 1D (Figure 2F and G). Furthermore, after performing the modified methods, the germination rates and pollen tube lengths did not decrease compared with those of the control (Figure 2H and I). These results indicate that plasmid spraying and vacuum conditions are helpful to enhance the delivery of gene modification components and do not effect pollen germination or growth. The transgenic pollen process used in this study took approximately 10 min (Figure 1D). Additionally, the main equipment used in these experiments were a NFJI device and a vacuum pump, both of which are economically accessible, with market prices of $230 and $110, respectively. Additionally, both the NFJI device and vacuum pump are portable, which is convenient for transgenic operations in the field. On the day of pear blossoming (April 10th), we introduced the apple gene MdGH3.1, which induces an early response to auxin, into pear pollen in the field for use in pollination. When the fruit matured (September 10th), the pear seeds were collected for qRT–PCR analysis. The results revealed that the expression level of the MdGH3.1 gene in the transgenic seeds was significantly greater than that in the wild-type seeds (Supplementary table S1).

**Figure 1.**
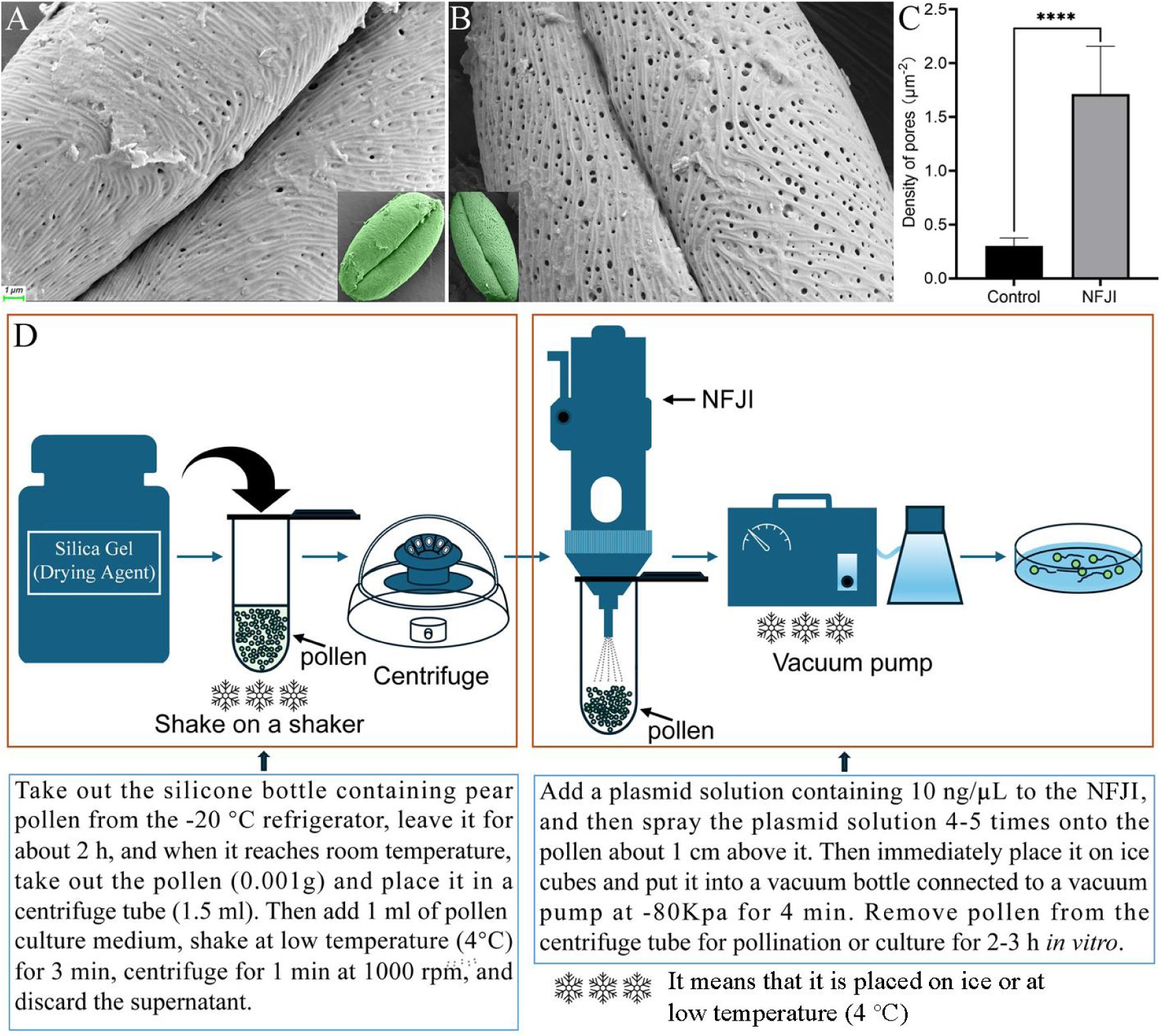
NFJI treatment of pollen. (A) Scanning electron microscopy (SEM) images of the pear pollen (‘Akitsuki’ cultivar). The inner image is the whole grain. (B) Distilled water was sprayed onto ‘Akitsuki’ pollen via NFJI. The inner image is the whole grain. (C) The number of micropores on the surface of the pollen. Control indicates pollen that has not been sprayed with distilled water; Spray indicates pollen that has been sprayed with distilled water. **** indicates *P* < 0.0001 (unpaired t test). (D) Process of genes transformation into pollen using NFJI and a vacuum pump.

**Figure 2.**
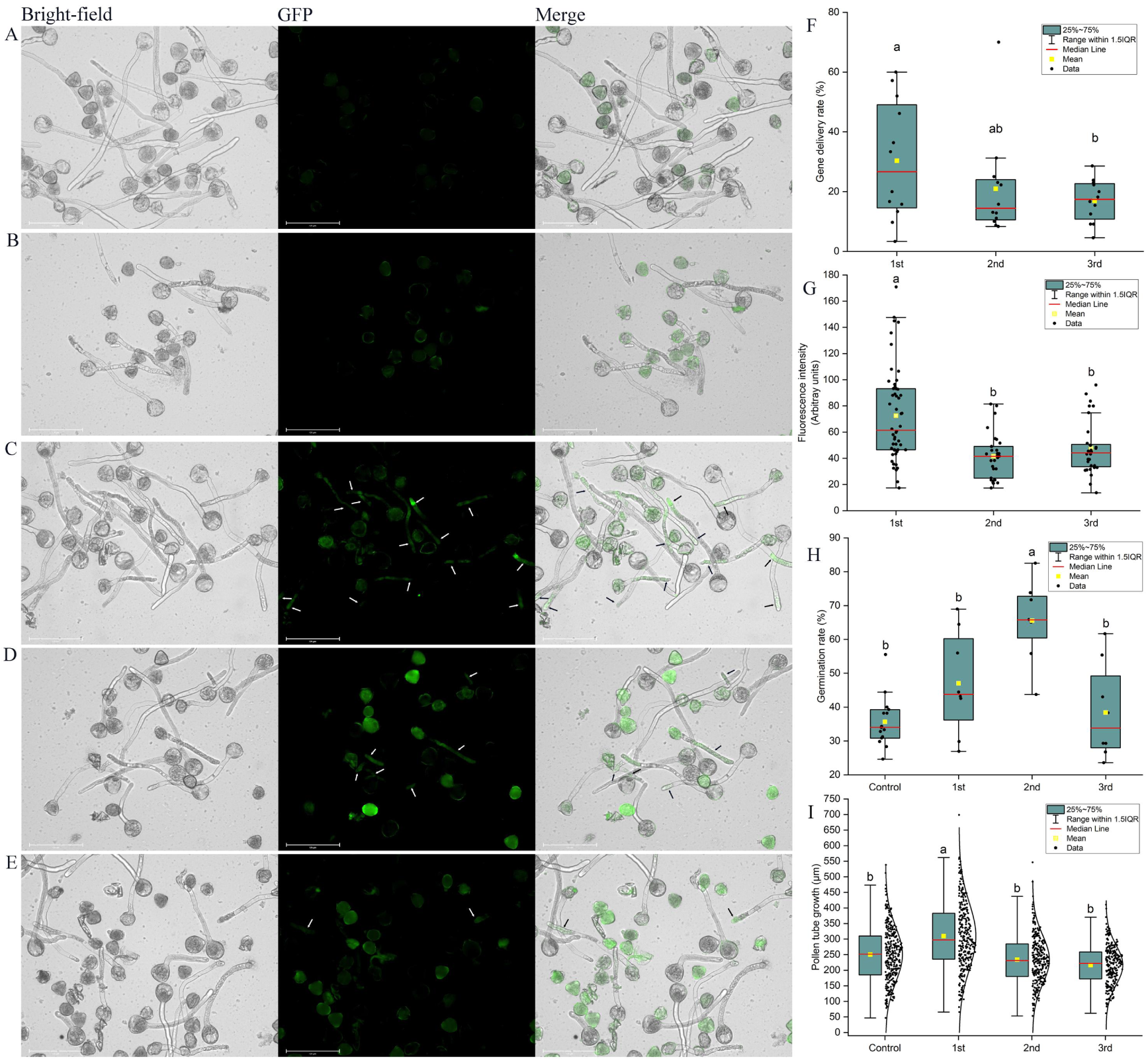
Verification of that the pollen was transformed with the GFP gene. Pollen culture was performed for 2 h at 25°C. (A) Nontransgenic ‘Akitsuki’ pollen (control). (B) ‘Akitsuki’ pollen was immersed in pollen culture medium containing the GFP plasmid (10 ng/μL), and then vacuum infiltration (-80 kPa) was performed. (C) ‘Akitsuki’ pollen was transformed with the GFP gene following the process shown in Figure 1D. The white arrow indicates where the fluorescent protein-encoding gene is expressed. (D) First, NFJI was used to spray distilled water onto ‘Akitsuki’ pollen, the pollen was placed in pollen culture medium containing the GFP plasmid (10 ng/μL), and vacuum infiltration was immediately performed for 4 min at -80 kPa. The white arrow indicates where the fluorescent protein-encoding gene is expressed. (E) Pollen culture medium containing the GFP plasmid (10 ng/μL) was sprayed onto ‘Akitsuki’ pollen using NFJI, and the pollen was cultivated directly without vacuum infiltration. The white arrow indicates where the fluorescent protein-encoding gene is expressed. (F) Gene delivery rates after different treatments. The x-axis indicates 1st, 2nd, and 3rd, which correspond to the treatment methods in (C), (D), and (E), respectively. Different lowercase letters indicate significant differences at *P* < 0.05. (G) Pollen tube fluorescence intensity after different treatments. The x-axis indicates 1st, 2nd, and 3rd, which correspond to the treatment methods in (C), (D), and (E), respectively. Different lowercase letters indicate significant differences at *P* < 0.05. (H) Germination rates of pollen after different treatments. The x-axis indicates Control, 1st, 2nd, and 3rd, which correspond to the treatment methods in (A), (C), (D), and (E). Different lowercase letters indicate significant differences at *P* < 0.05. (I) Growth of pollen tubes after different treatments. The x-axis indicates Control, 1st, 2nd, and 3rd, which correspond to the treatment methods in (A), (C), (D), and (E). Different lowercase letters indicate significant differences at *P* < 0.05.

### Validation of gene function using tobacco as a model plant

Tobacco plants have a flowering to seed setting cycle that is shorter than that of pear plants. Additionally, tobacco plants can reproduce multiple times within one year under suitable growth conditions. In addition, model plants can be used to study of specific biological phenomena or biotechnologies (Prusinkiewicz, 2004; Ray et al., 2022). Using the transgenic process shown in Figure 1D, two genes (GFP and MdGH3.1) were delivered into tobacco pollen separately, and after pollination was performed, the mature seeds were collected for sowing. Tobacco pollen transformed with the GFP gene exhibited fluorescent pollen tubes after cultivation (Supplementary figure S4). Given the autofluorescence of chlorophyll when observed under a fluorescence microscope, which can interfere with the detection of specific fluorescent proteins (Donaldson, 2020), a fluorescence microscope was used in this study to observe the tobacco roots and root hairs after GFP gene transfer. Although the roots also contain lignin and flavonoids, which may also interfere with the detection of specific fluorescent proteins (Wang et al., 2023), the fluorescence intensity of the wild-type roots was significantly lower than that of the transgenic roots. In particular, the root hairs of the wild-type tobacco did not fluoresce, whereas the transgenic tobacco root hairs exhibited considerable fluorescence (Figure 3).

**Figure 3.**
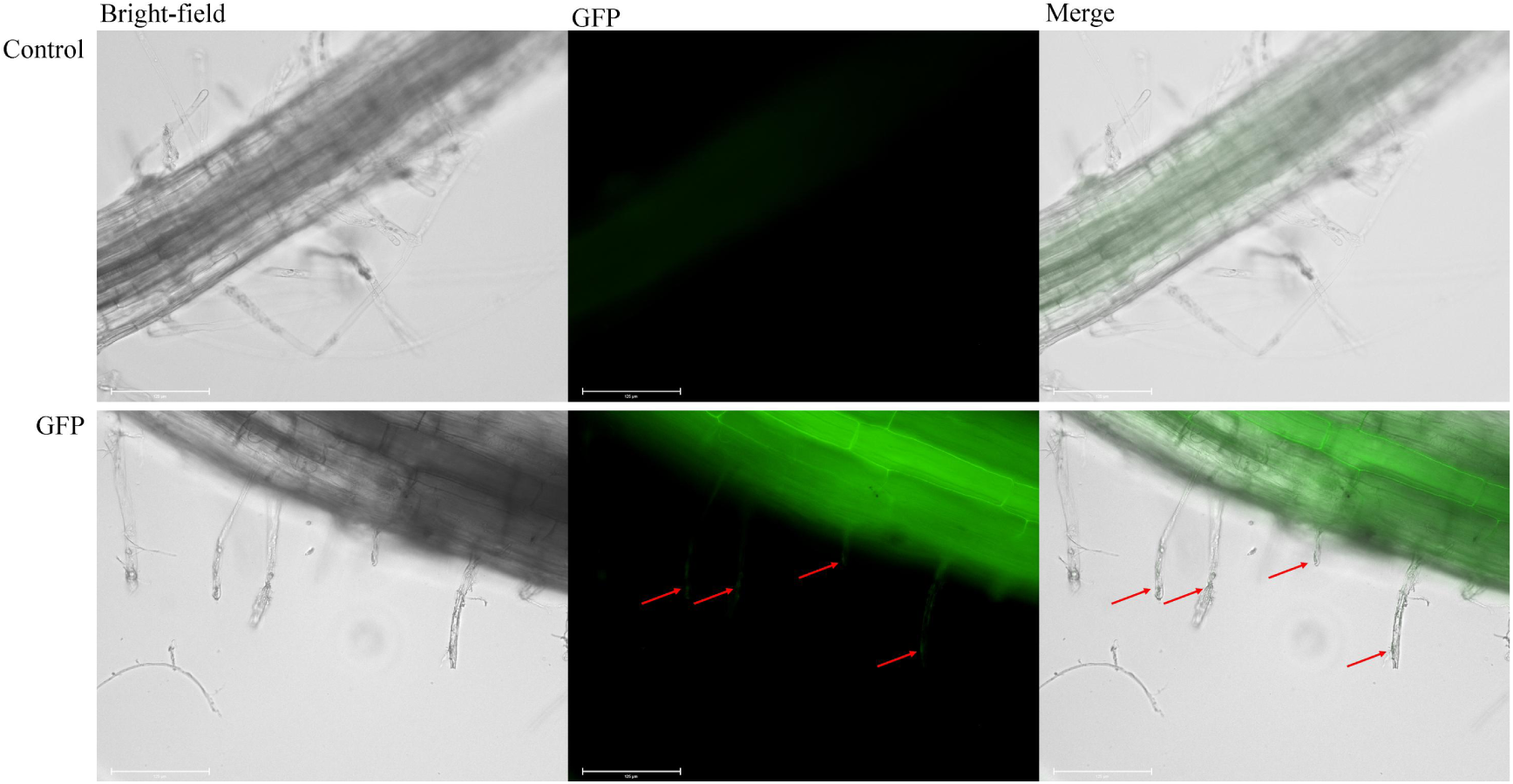
Using tobacco as a model plant. The GFP reporter gene was introduced into tobacco using the method shown in Figure 1D, and the GFP reporter gene was expressed in the roots and root hairs.

We randomly selected 21 tobacco seedlings transformed with the MdGH3.1 gene for quantitative detection, among which 15 overexpressed MdGH3.1 (indicated by the colored cells in Supplementary table S2), demonstrating that the transformation success rate of this method can reach 71%. The GH3.1 gene product enables inactivates auxin by converting it into inactive forms such as IAA-ASP and IAA-Glu (Hayashi et al., 2021), which thereby inhibits root hair growth (Breakspear et al., 2014; Wang et al., 2023). Compared with wild-type plants, tobacco plants transformed with the MdGH3.1 gene presented dwarfism (Figure 4A), longer lateral roots but fewer, shorter root hairs (Figure 4B-E). We divided these 15 seedlings into two types: those whose relative expression level was greater than 2 but lower than 10 were classified as the low-overexpression type (blue cells in Supplementary table S2), and those whose relative expression level was greater 20 were classified as the high-overexpression type (green cells in Supplementary table S2).

**Figure 4.**
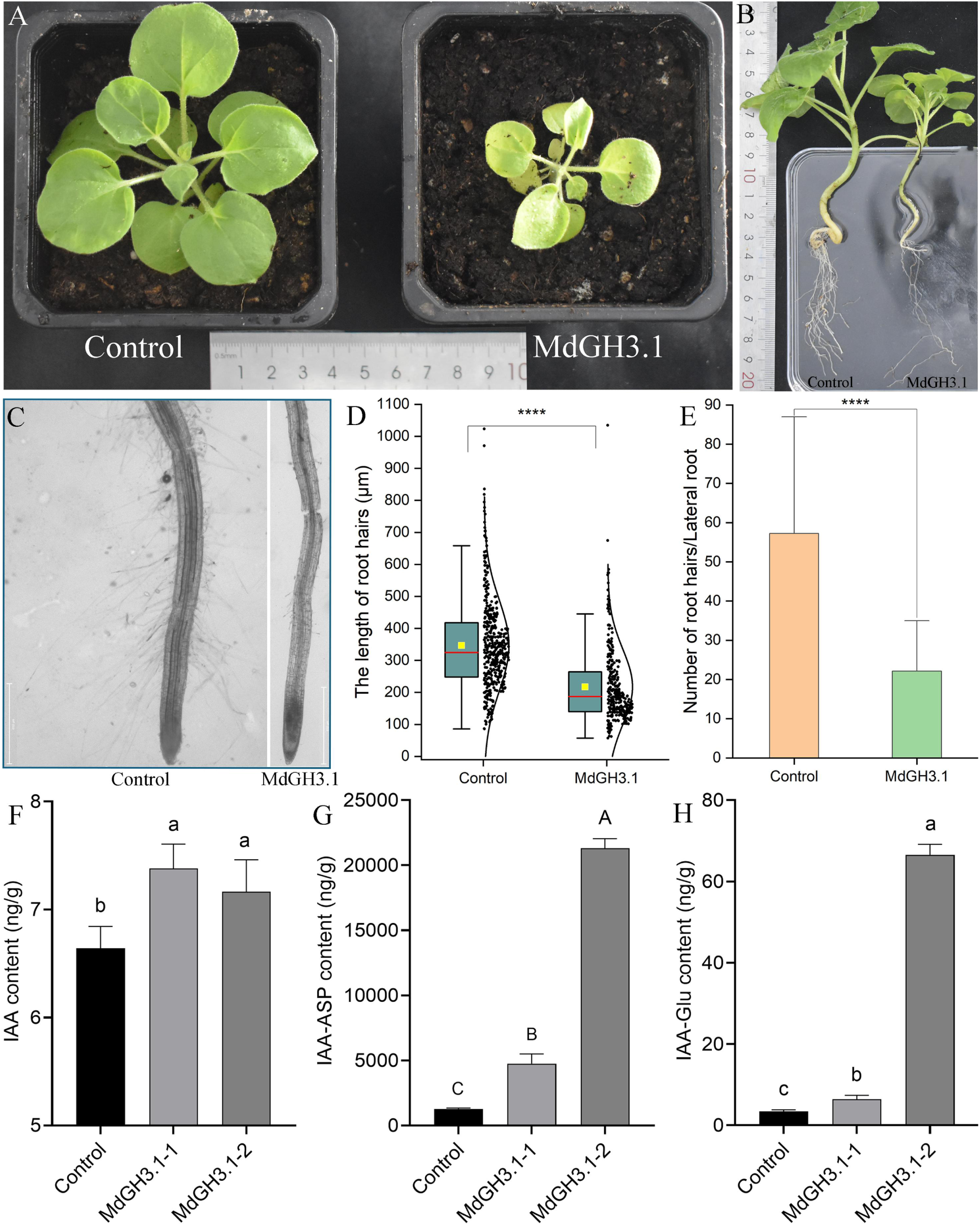
The apple gene MdGH3.1 was introduced into tobacco using the method shown in Figure 1D. (A) Comparison of the sizes of the transgenic and control (wild-type) tobacco. (B) Comparison of the root systems of transgenic and control (wild-type) tobacco. (C) Comparison of the root hairs of transgenic and control (wild-type) tobacco. (D) The root hair lengths of 3 control tobacco plants (458 root hairs) and 7 transgenic tobacco plants (338 root hairs) were measured. **** indicates *P* < 0.0001 (unpaired t test). (E) The number of root hairs on 8 roots of 3 control tobacco plants and 14 roots of 7 transgenic tobacco plants was determined. **** indicates *P* < 0.0001 (unpaired t test). (F) Comparison of the auxin contents in tobacco plants with different MdGH3.1 gene expression levels. According to Supplementary table S2, tobacco plants with a relative expression level of the MdGH3.1 gene greater than 2 but less than 10 were classified as MdGH3.1-1, and those with a relative expression level greater than 20 were classified as MdGH3.1-2. Different lowercase letters indicate significant differences at *P* < 0.05. (G) Comparison of IAA-ASP contents in tobacco plants with different MdGH3.1 gene expression levels. Different capital letters indicate significant differences at *P* < 0.01. (H) Comparison of the IAA-Glu contents in tobacco plants with different MdGH3.1 gene expression levels. Different lowercase letters indicate significant differences at *P* < 0.05.

Additionally, the auxin contents in the two types of MdGH3.1 overexpressing tobacco were greater than that in wild-type tobacco. Although the auxin content in the low-overexpression tobacco plants was greater than that in the high-overexpression tobacco plants, the difference was not significant (Figure 4F). However, the contents of IAA-ASP and IAA-Glu in tobacco plants overexpressing MdGH3.1 were significantly greater than those in wild-type tobacco (Figure 4G and H), and the high-overexpression tobacco plants also had significantly greater contents IAA-ASP and IAA-Glu than the low-overexpression plants (Figure 4G and H). In addition, the GH3 family of genes are all auxin-responsive genes, among which GH3.1 mainly converts auxin into IAA-ASP rather than IAA-Glu (Wang et al., 2023); this is also consistent with the results of this study (Figure 4G and H). Therefore, NFJI can effectively deliver exogenous genes into pollen for their expression in offspring after sexual reproduction.

### Applications to pollen of other plants

The GFP gene was successfully introduced into the pollen of dicotyledonous plants such as apples (Figure 5A) and tomatoes using a NFJI, resulting in their pollen tubes exhibiting significant fluorescence (Supplementary figure S5). A significant challenge in the application of Agrobacterium-mediated transformation technology to monocotyledonous plants is the low infection efficiency (Rahman et al., 2024). We successfully introduced the GFP gene into the pollen of the monocotyledonous plant maize using NFJI (Figure 5B). MdGH3.1 was subsequently transformed into another model plant, *A. thaliana*, of the Brassicaceae family, and 14 Arabidopsis seedlings were randomly selected for gene expression analysis. The results revealed that 11 seedlings highly expressed MdGH3.1, indicating a transformation rate of 79% (Supplementary table S3). Compared with that of wild-type plants, the phenotype of these transgenic *A. thaliana* was similar to that of the transgenic tobacco, as the plants were smaller with longer roots but fewer, shorter root hairs (figure 5C-E).

**Figure 5.**
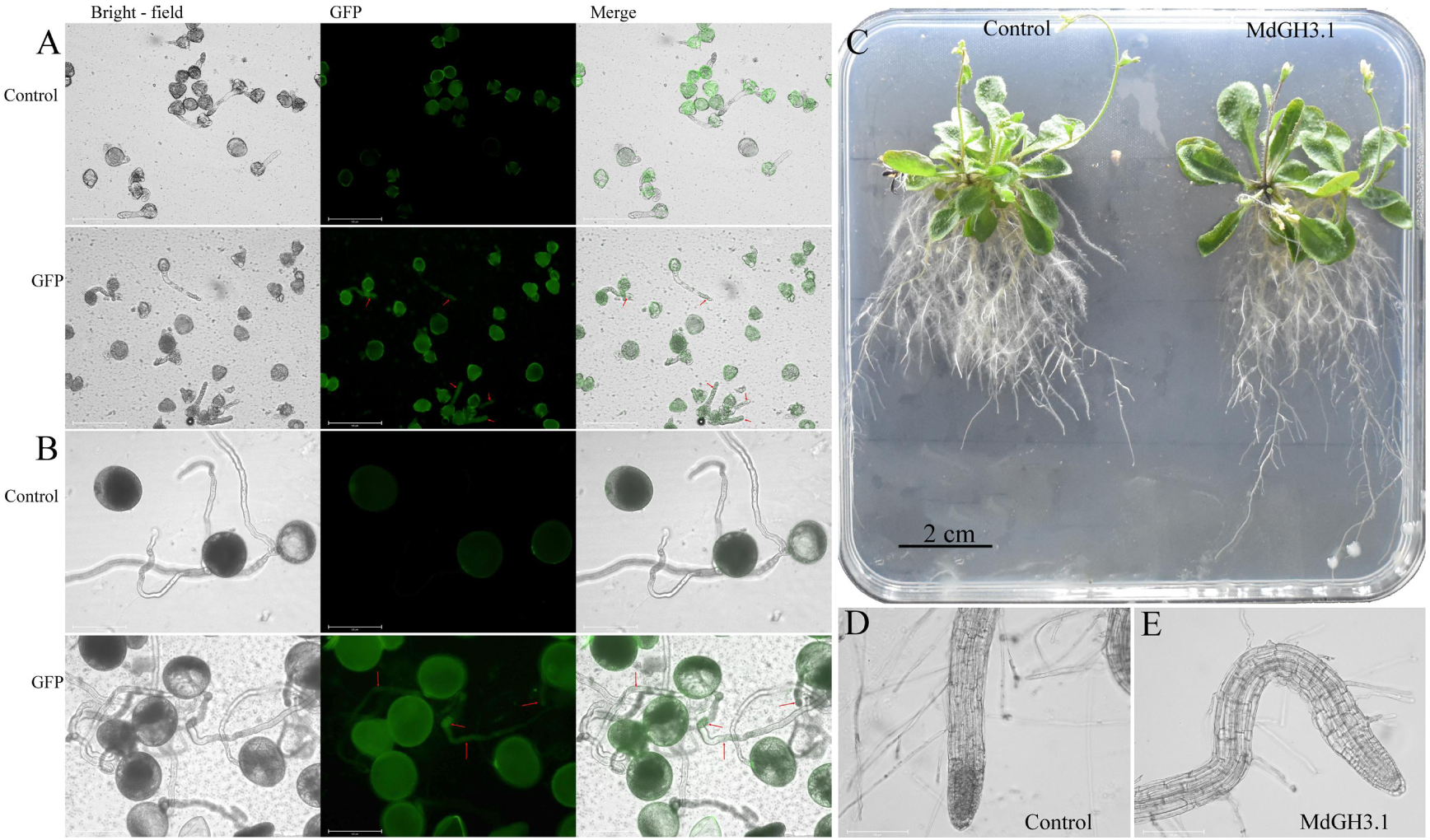
Application of this transgenic method to other species. (A) Transgenic apple pollen; the control was nontransgenic ‘Fuji’ apple pollen. GFP indicates that the GFP gene was transferred into apple pollen using the procedure outlined in Figure 1D. (B) Transgenic corn pollen; the control was nontransgenic corn pollen. GFP indicates that the GFP gene was transferred into corn pollen using the procedure outlined in Figure 1D. (C) The apple gene MdGH3.1 was transformed into the model plant Arabidopsis using the method outlined in Figure 1D, and the sizes and root systems of the transgenic and control (wild-type) Arabidopsis plants were compared. (D) Root hairs of control (wild-type) Arabidopsis. (E) Root hairs of Arabidopsis transformed with the MdGH3.1 gene.

## Discussion

Given that plants have cell walls, directly introducing exogenous genetic material into the cell interior poses significant challenges (Furuhata et al., 2019). To solve this problem, researchers have adopted a variety of methods, including biological, chemical, and physical methods. Among them, physical approaches such as gene gun technology, microinjection, and electroporation are increasingly favored by researchers for their ability to induce efficient cell wall penetration (Rivera et al., 2014). Nanomagnetic technology and biotransduction are cutting-edge physical conversion strategies, and their current research is focused primarily on the efficiency of transient transformation (Khan et al., 2022; Tawade and Wasewar, 2023; Wu et al., 2024). Moreover, these technologies rely on tissue regeneration techniques, similar to Agrobacterium-mediated methods. However, there are substantial difficulties with tissue culture and regeneration of many plant species (Cunningham et al., 2018). Therefore, transgenic techniques that do not require tissue culture have attracted attention because of their high transformation efficiency and short transformation cycles ^11,36^. The NFJI and vacuum infiltration methods used in this study are both physical transgenic methods that can efficiently introduce exogenous genes into pollen grains. As important carriers in the transformation process, pollen grains have great research value (Park et al., 2024). As long as a gene can be transferred into pollen, it can be stably introduced into the next generation through pollination and fertilization (Eapen, 2011). Nanomagnetic technology has been applied to transfer exogenous genes into maize and cotton through their pollen, and these genes were stably expressed in the next generation. In this study, we introduced the MdGH3.1 gene from apples (Rosaceae) into the pollen of tobacco (Solanaceae) and Arabidopsis (Brassicaceae), which was expressed in the next generation after pollination, and verified that it was functional. Generating transgenic plants via pollination eliminates the need for time-consuming and labor-intensive tissue culture. Nanomagnetic technology has been employed to successfully transfer exogenous genes into maize pollen. This technique is based on the maize pollen having germination pores with diameters about 6 µm (Wang et al., 2022). NFJI induces the formation of more micropores on the pollen surface, providing channels for the introduction of exogenous genes. Moreover, a vacuum environment facilitates the intracellular transfer of exogenous genes. Therefore, after treating pollen via NFJI, vacuum infiltration technology can be further applied to increase the efficiency of exogenous gene transfer.

Pollen dormancy is crucial for utilizing pollen as a genetic resource in plant breeding ^39^. During the double fertilization of plants, it is essential that the pollen remains dormant throughout the dissemination and preservation stages (Firon et al., 2012). Pollen can initiate germination and pollen tube growth only after contacting the female stigma, thus achieving successful double fertilization. Pollen that has undergone specific treatments does not remain dormant when cultured in liquid medium ^41^. Studies have shown that various types of pollen can begin to germinate within 30 min in medium, and once it has germinated, it is no longer suitable for subsequent pollination processes. The genes in this study were delivered via NFJI technology, which is characterized by a short operation time and high efficiency, and the entire process was completed in approximately 10 min. Importantly, shortening the transgenic treatment duration has great advantages for improving pollination efficiency.

Pollen viability refers to the ability of pollen to survive under different environmental conditions and to maintain its abilities to germinate form pollen tube tips on compatible stigmas, which are key elements in plant reproductive processes (Althiab-Almasaud et al., 2024; Edlund et al., 2004; Pacini and Dolferus, 2019). The aim of this study was to facilitate genetic transformation by inducing the formation of more micropores on the pollen surface via NFJI. In this research, nanotungsten powder was sprayed on the pollen during NFJI. However, this method led to significant reductions in the pear pollen germination rate and pollen tube growth, thus negatively affecting pollen viability (Supplementary figure S6). Therefore, this method was abandoned.

In summary, the use of pollen as a vector for genetic transformation is an in vivo transformation technique that does not rely on a specific genotype and is applicable to both dicotyledonous and monocotyledonous plants (such as maize). This method can efficiently introduce exogenous genes into the pollen of various families and genera of plants. The entire transformation process is efficient (taking approximately 10 min), easy to apply, and requires affordable, portable equipment, making field application possible. Therefore, this technology has broad application potential in a variety of plant species.

## Materials and methods

### Materials

#### Pollen collection

In this study, we selected ‘Akitsuki’ pears (*Pyrus pyrifolia* Akitsuki) and ‘Fuji’ apples (*Malus domestica* Borkh. CV. Fuji), which are widely cultivated in China. Pollen samples were collected in April from the Shandong Laixi Breeding Farm (China). Tobacco (*N. benthamiana*), *Arabidopsis thaliana* (*A. thaliana* (L.) Heynh.), and tomato (*Solanum lycopersicum*) pollen samples were provided by the laboratory. Additionally, the maize (*Zea mays ssp*. mays) pollen samples were kindly donated by Chinese Academy of Tropical Agricultural Sciences.

### Culture medium

Pear and apple pollen culture media was composed of 0.55 mmol/l Ca(NO_3_)_2_, 1.6 mmol/l H_3_BO_3_, 1.6 mmol/l MgSO_4_, 1 mmol/l KNO_3_, 440 mmol/l sucrose and 5 mmol/l 5,2-(N-morpholino) ethane sulfonic acid hydrate (MES); the pH was adjusted to 6.0 with Tris.

Tomato pollen culture media was composed of 12% sucrose, 1.94 mM H_3_BO_3_, 11.55 μM GA_3_, and 1.48 μM vitamin B1.

Maize pollen media was composed of 20% sucrose, 808.67 μM H_3_BO_3_, 2.703 μM CaCl_2_, 2.1 μM MgCl_2_, 989.12 μM KNO_3_, and 101.05 μM GA_3_.

Tobacco pollen culture media was composed of 15% sucrose, 24.27 μM H_3_BO_3_, and 54 μM CaCl_2_.

Arabidopsis seed culture media was composed of 2.47 g/l 1/2 MS medium (without agar or sucrose), 30 g/l sucrose, and and 7 g/l agar (pH 5.8). The samples were first vernalized at 4°C for 24 h and then grown in a 22°C incubator for 25 d.

### Methods

#### Electron microscopy analysis of the pollen samples

After being sprayed with gold, the pollen grains were observed under a scanning electron microscope (ZEISS Merlin Compact, Germany) at 3.0 kV.

#### Plasmids

The 35S::GFP plasmid was a generous gift from the State Key Laboratory of Crop Genetics and Germplasm Enhancement at Nanjing Agricultural University. To create the 35S::MdGH3.1 plasmid, GFP in the 35S::GFP plasmid was replaced with MdGH3.1.

#### NFJI operation

1. Tighten the nozzle to the corresponding interface of the cone; then, fit the glass tube onto the cone.
2. Use a pipette to add 1 ml plasmid solution (10 ng/µL) into a glass tube. Then, insert the sleeve into the glass tube and lock it to the cone by rotation.
3. Pull the pressure handle until a click is heard to complete pressurization. Then, return the pressure handle to its original position with the nozzle facing downward, and press the operation button to complete one injection.
4. Repeat the above 3 steps, remove the air, and if liquid is being expelled, formal spraying treatment can be performed. During spraying, the nozzle should be 0.5–1 cm from the pollen grain.
5. After each application of the plasmid mixture in the glass tube, use the matching tool to unscrew the hexagonal pressure cap inside the conical body in the opposite direction, replace the rubber ring under the metal flat pad with a cleaned (sterilized) new rubber ring, and restore it to the original state with the matching tool for later use.

#### Fluorescence detection

After 2-3 h of culture, the transgenic pollen in the tube was observed under a fluorescence microscope (EVOS Auto 2, Thermo Fisher, USA) with the selected GFP light cube (excitation of 488 nm). Additionally, the leaves of the transgenic seedlings were observed via stereo-fluorescence microscopy (SZX12, Olympus, USA).

### Determination of gene expression via quantitative real-time RT-PCR(qRT– PCR)

qRT‒PCR was performed to estimate the mRNA expression levels. Total RNA was extracted from the pollen and styles with TRIzol reagent (Invitrogen), and first-strand cDNA was synthesized from 2 μg of total RNA using an Omniscript RT kit (Qiagen). The primers used for MdGH3.1 were F: CTCGGATATGGCCAAACACG and R: AGAAGGCTTGCACATAGGGT. Gene expression data were normalized to actin gene expression and are presented as a ratio compared to the relevant control (assigned a value of 1).

### Pollen tube-mediated transformation and calculation of pollen transformation efficiency

The pollen tube transformation method was performed according to the process shown in Figure 1D for pollen genetic transformation. GFP transgenic pollen was cultivated for 2 h, and then, 1 μL of the sample was placed on a slide for observation under a fluorescence microscope. Additionally, the gene delivery rate (%) was calculated as the ratio of fluorescent pollen tubes to the total number of pollen tubes.

### Root hair counting and length measurements

The roots of the tobacco and Arabidopsis plants were removed and washed with water, after which microscopic photos of the root hair zone were taken. Then, ImageJ was used to count the number of root hairs and measure their length.

### Statistical analysis

The data are presented as the means ± standard deviations (SDs) and were analyzed with GraphPad Prism version 7.0 software (GraphPad Software, Inc., La Jolla, CA, USA) and Origin 8.0 (OriginLab, USA). The significance of the differences between experimental groups were assessed using Student’s t test.

## Acknowledgments

We thank Professor Juyou Wu of Nanjing Agricultural University for providing the GFP plasmid. Thank you to Researcher Xiaowei Ma from the Chinese Academy of Tropical Agricultural Sciences for providing the maize pollen.

## Author contributions

Q.H.Y. conceived and designed the study. W.H, Q.W, performed the experiments. Q.H.Y. analyzed the data. Z.X collected pollen. Q.H.Y. wrote the manuscript text and prepared Figures 1-5. All authors reviewed the manuscript.

## Declarations

The authors declare that they have no conflicts of interest regarding the content of this article.

## Data Availability declaration

All relevant data are within the manuscript and its Additional files.

## Ethical approval

Ethics approval was not required for this research.

