## Supplementary figures and tables for "Simple, fast, low-cost method to produce transgenic pollen"

### Supplementary figures information

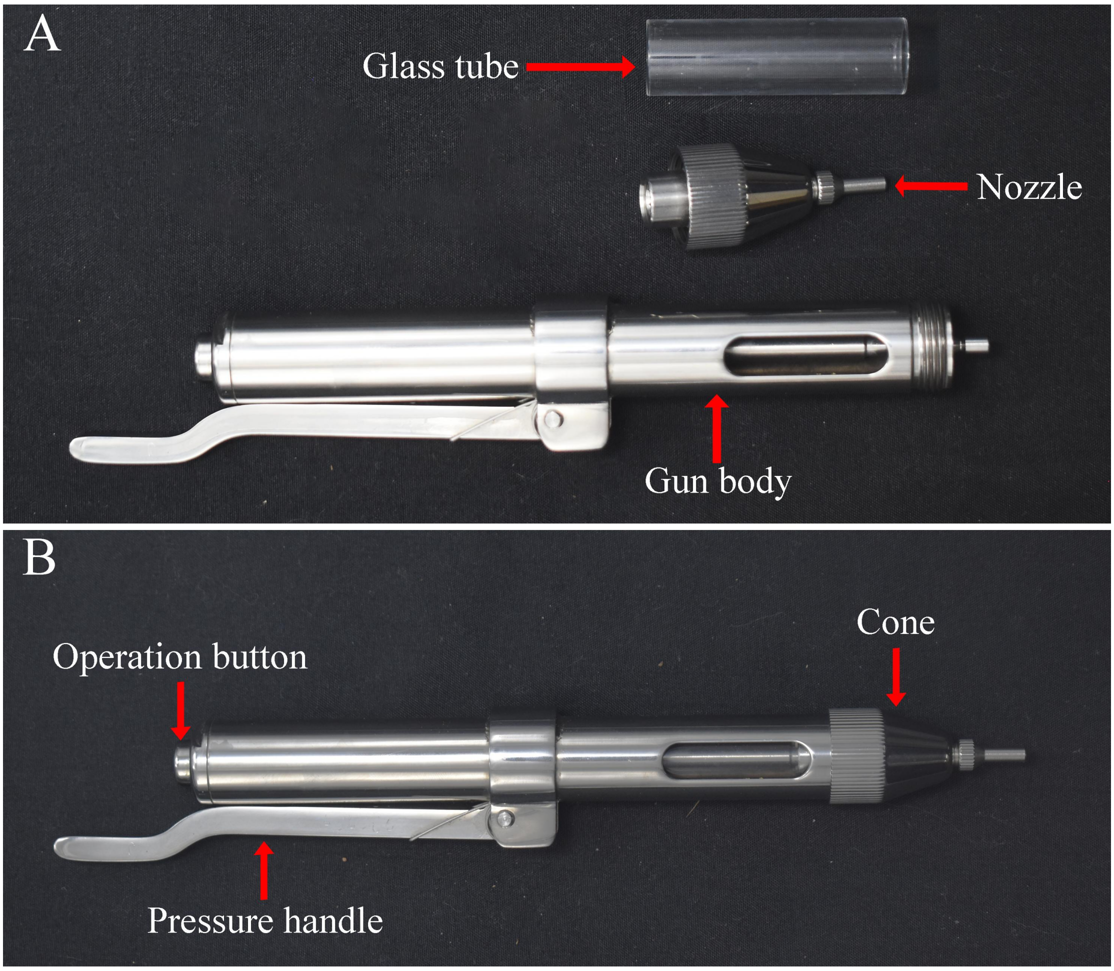

**Supplementary figure S1. Needle-free jet injection (NFJI).** (A) The main NFJI accessories. (B) An assembled NFJI device.

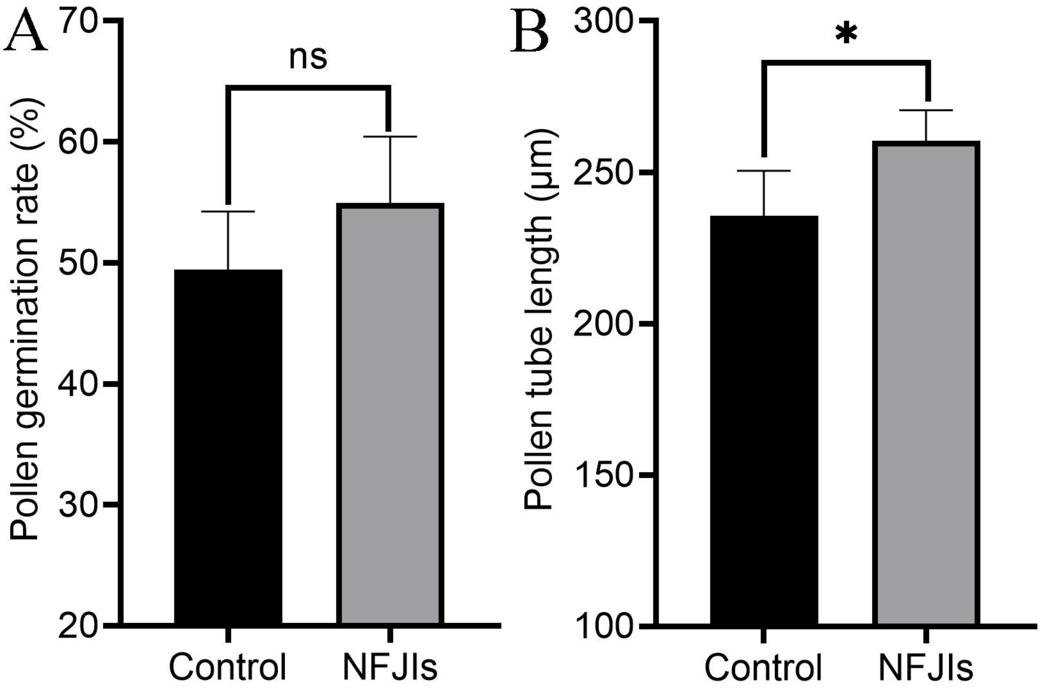

**Supplementary figure S2. 'Akitsuki' pollen germination rate (A) and pollen tube growth (B).** Control indicates pollen that has not been sprayed with distilled water, and NFJI indicates pollen that was sprayed with distilled water via NFJI.

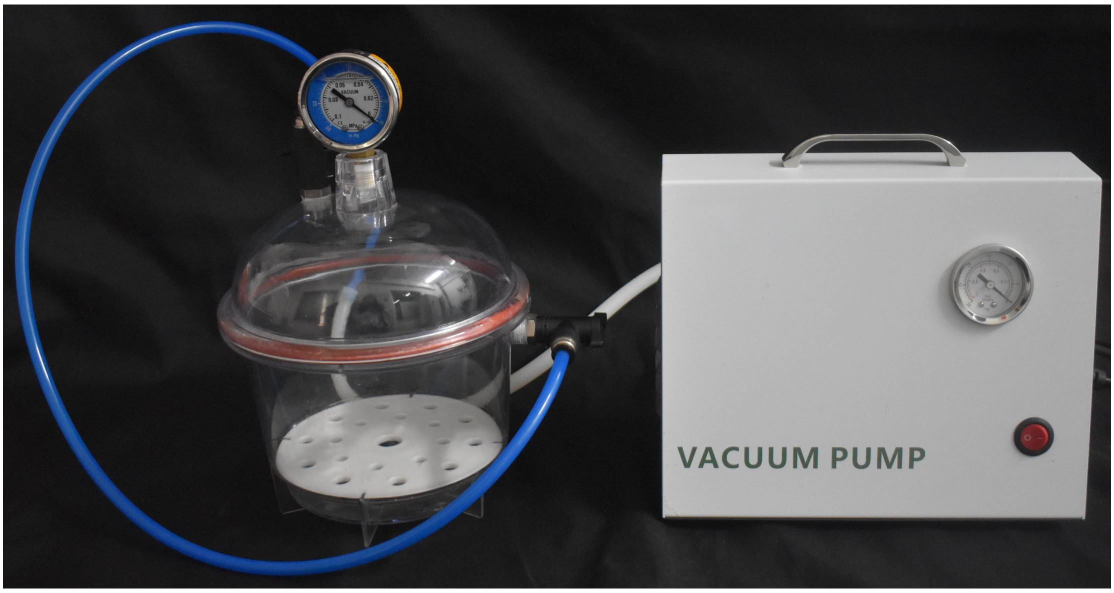

**Supplementary figure S3. An easy to carry vacuum pump was used.** After the pollen was placed ino the vacuum bottle, the vacuum pump was turned on, and upon reaching a vacuum of -80 Kpa, a timer was set for 4 min, after which time the vacuum pump was turned off immediately.

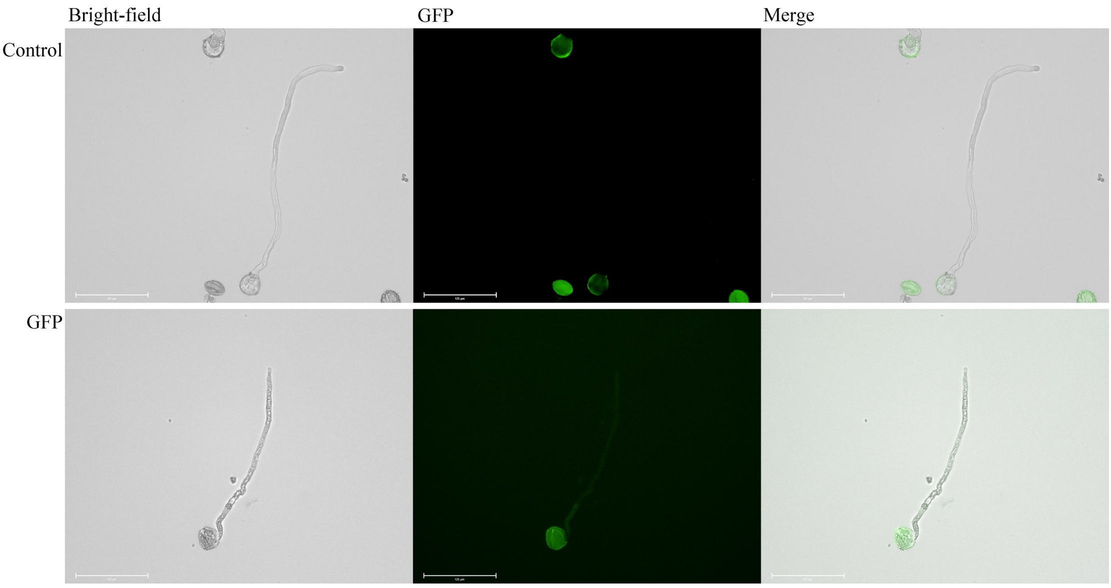

**Supplementary figure S4.** Tobacco pollen was transformed with the GFP gene following the process shown in Figure 1D. Nontransgenic pollen was used as a control.

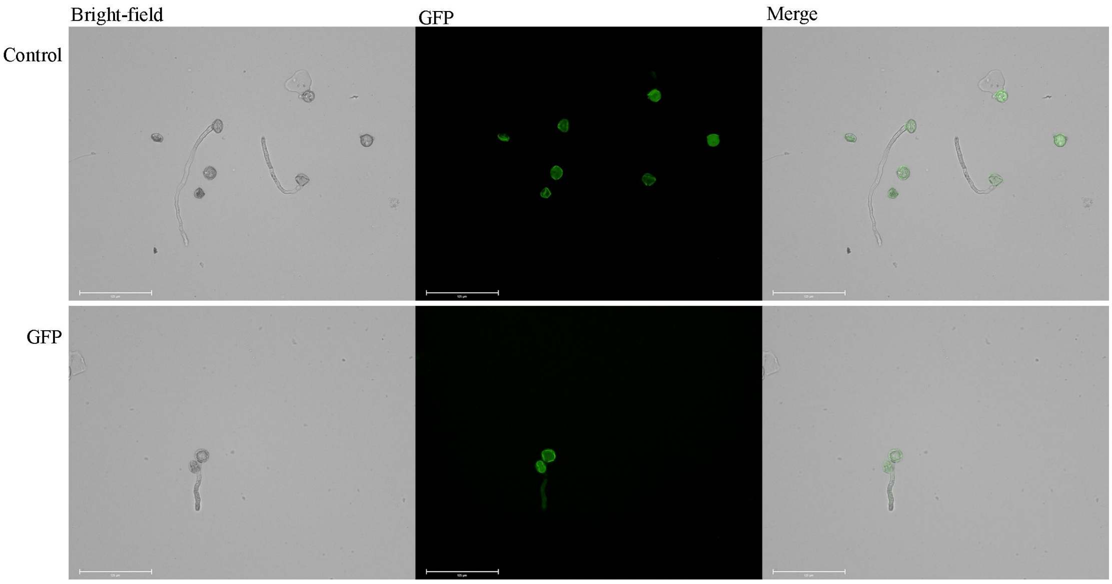

**Supplementary figure S5.** Tomato pollen was transformed with the GFP gene following the process shown in Figure 1D. Nontransgenic pollen was used as a control.

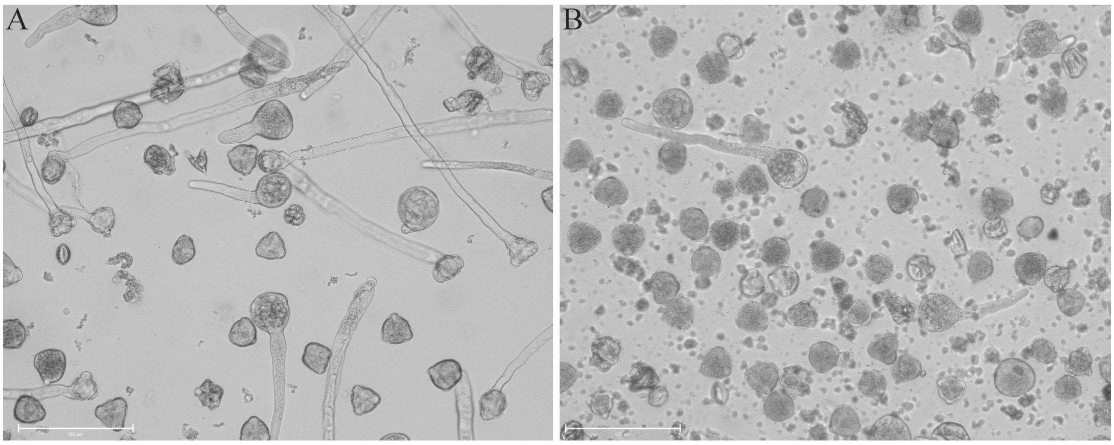

**Supplementary figure S6.** Nanoscale tungsten powder was incorporated into NFJI and sprayed onto ‘Akitsuki’ pollen. (A) Control. Pollen that was not sprayed. (B) Pollen sprayed with nanoscale tungsten powder.

**Supplementary tables information**

**Supplementary table S1.** On April 10, 2024, the MdGH3.1 gene was transferred into 'Akitsuki' pear pollen in the field using the method shown in Figure 1D and then used to pollinate 'Xueqing' pears (compatible pollination). Nontransgenic 'Akitsuki' pear pollen was used to pollinate 'Xueqing' pears as a control. On September 10th, mature 'Xueqing' pear fruits were harvested, and the seeds from three randomly selected control pear fruits and three genetically modified pear fruits were collected. Quantitative analysis of MdGH3.1 gene expression in the seeds. Two out of three genetically modified pear seeds highly expressed the MdGH3.1 gene.

**Supplementary table S2.** Quantitative analysis of the relative expression levels of MdGH3.1 in transgenic tobacco. The control is wild-type tobacco. The color of the cells indicates the extent of MdGH3.1 overexpression. Blue cells indicate relative expression levels between 2 and 10, considered low overexpression. Green cells indicate relative expression levels greater than 20, considered high overexpression. Yellow cells indicate MdGH3.1 overexpression.

**Supplementary table S3.** Quantitative analysis of the relative expression levels of MdGH3.1 in transgenic *Arabidopsis thaliana*. The control is wild-type *Arabidopsis thaliana*. Yellow cells indicate MdGH3.1 overexpression.

| Sample | cp | Actin | Actin (Average) | cp1 | cp2 | △cp | Relative gene expression | Relative gene expression (Average) |
|---|---|---|---|---|---|---|---|---|
| Control-1 | 28.74 | 26.13 | 26.065 | 2.675 | 3.4925 | -0.8175 | 1.762349426 | 1.068372131 |
| | 29.6 | 26.11 | 26.065 | 3.535 | 3.4925 | 0.0425 | 0.970970924 | |
| | 30.12 | 26.12 | 26.065 | 4.055 | 3.4925 | 0.5625 | 0.677127773 | |
| | 29.77 | 25.9 | 26.065 | 3.705 | 3.4925 | 0.2125 | 0.8630404 | |
| MdGH3.1-1 | 26.02 | 25.71 | 25.7825 | 0.2375 | 3.4925 | -3.255 | 9.546685944 | 14.65299596 |
| | 25.11 | 25.59 | 25.7825 | -0.6725 | 3.4925 | -4.165 | 17.93865725 | |
| | 25.33 | 25.87 | 25.7825 | -0.4525 | 3.4925 | -3.945 | 15.40151109 | |
| | 25.3 | 25.96 | 25.7825 | -0.4825 | 3.4925 | -3.975 | 15.72512958 | |
| Control-2 | 27.11 | 22.55 | 22.03 | 5.08 | 4.285 | 0.795 | 0.576343173 | 1.052369609 |
| | 26.2 | 22.34 | 22.03 | 4.17 | 4.285 | -0.115 | 1.082975046 | |
| | 26.15 | 21.93 | 22.03 | 4.12 | 4.285 | -0.165 | 1.121166078 | |
| | 25.8 | 21.3 | 22.03 | 3.77 | 4.285 | -0.515 | 1.42899414 | |
| MdGH3.1-2 | 29.31 | 30.23 | 30.0775 | -0.7675 | 4.285 | -5.0525 | 33.18593461 | 36.13130082 |
| | 29.43 | 30.56 | 30.0775 | -0.6475 | 4.285 | -4.9325 | 30.5372872 | |
| | 28.84 | 29.52 | 30.0775 | -1.2375 | 4.285 | -5.5225 | 45.96615223 | |
| | 29.24 | 30 | 30.0775 | -0.8375 | 4.285 | -5.1225 | 34.83582922 | |
| Control-3 | 27.57 | 24.8 | 24.28 | 3.29 | 2.745 | 0.545 | 0.685391402 | 1.11154321 |
| | 27.71 | 24.83 | 24.28 | 3.43 | 2.745 | 0.685 | 0.622005827 | |
| | 26.73 | 24.7 | 24.28 | 2.45 | 2.745 | -0.295 | 1.226884977 | |
| | 26.09 | 22.79 | 24.28 | 1.81 | 2.745 | -0.935 | 1.911890635 | |
| MdGH3.1-3 | 30.53 | 24.51 | 23.8175 | 6.7125 | 2.745 | 3.9675 | 0.063923934 | 0.112191254 |
| | 29.7 | 23.93 | 23.8175 | 5.8825 | 2.745 | 3.1375 | 0.113636641 | |
| | 29.18 | 23.8 | 23.8175 | 5.3625 | 2.745 | 2.6175 | 0.162949858 | |
| | 29.77 | 23.03 | 23.8175 | 5.9525 | 2.745 | 3.2075 | 0.108254582 | |

**Supplementary table S1.**On April 10, 2024, the MdGH3.1 gene was transferred into ‘ Akitsuki ’ pear pollen in the field using the method shown in Figure 1D and then used to pollinate ‘ Xueqing ’ pears (compatible pollination). Nontransgenic ‘ Akitsuki ’ pear pollen was used to pollinate ‘ Xueqing ’ pears as a control. On September 10th, mature ‘ Xueqing ’ pear fruits were harvested, and the seedsfrom three randomly selected control pear fruits and three genetically modified pear fruits were collected. Quantitative analysis of MdGH3.1 gene expression in the seeds. Two out of three genetically modified pear seeds highly expressed the MdGH3.1 gene.

| Time | Gene | Sample | cp | Actin | Actin (Average | cp1 | cp2 | △cp | Relative gene expression | Relative gene expression (Average) |
|---|---|---|---|---|---|---|---|---|---|---|
| 24.10.23 | MdGH3.1 | Tobacco (Control) | 35.38 | 23.12 | 23.09666667 | 12.2833 | 9.38 | 2.90333333 | 0.1336625 | 1.874895373 |
| | | | 31.15 | 23.01 | 23.09666667 | 8.05333 | 9.38 | -1.3266667 | 2.508224819 | |
| | | | 30.9 | 23.16 | 23.09666667 | 7.80333 | 9.38 | -1.5766667 | 2.982798801 | |
| | | Tobacco -1 | 33.49 | 25.04 | 25.12666667 | 8.36333 | 9.38 | -1.0166667 | 2.023238881 | 1.774602657 |
| | | | 33.84 | 25.03 | 25.12666667 | 8.71333 | 9.38 | -0.6666667 | 1.587401052 | |
| | | | 33.73 | 25.31 | 25.12666667 | 8.60333 | 9.38 | -0.7766667 | 1.713168038 | |
| | | Tobacco-2 | 34.35 | 23.27 | 23.25 | 11.1 | 9.38 | 1.72 | 0.303548721 | 0.313538425 |
| | | | 34.5 | 23.13 | 23.25 | 11.25 | 9.38 | 1.87 | 0.273573425 | |
| | | | 34.09 | 23.35 | 23.25 | 10.84 | 9.38 | 1.46 | 0.363493129 | |
| | | Tobacco-3 | 34.02 | 21.69 | 21.74666667 | 12.2733 | 9.38 | 2.89333333 | 0.134592196 | 0.12604541 |
| | | | 34.35 | 21.72 | 21.74666667 | 12.6033 | 9.38 | 3.22333333 | 0.107073002 | |
| | | | 34 | 21.83 | 21.74666667 | 12.2533 | 9.38 | 2.87333333 | 0.136471033 | |
| 24.10.28 | MdGH3.1 | Sample | cp | Actin | Actin (Average | cp1 | cp2 | △cp | Relative gene expression | Relative gene expression (Average) |
| | | Tobacco (Control) | 32.81 | 23.5 | 23.52666667 | 9.28333 | 9.53333 | -0.25 | 1.189207115 | 1.02236848 |
| | | | 33.5 | 23.49 | 23.52666667 | 9.97333 | 9.53333 | 0.44 | 0.737134609 | |
| | | | 32.87 | 23.59 | 23.52666667 | 9.34333 | 9.53333 | -0.19 | 1.140763716 | |
| | | Tobacco-4 | 34.94 | 23.54 | 23.55333333 | 11.3867 | 9.53333 | 1.85333333 | 0.276752195 | 0.544492808 |
| | | | 33.75 | 23.59 | 23.55333333 | 10.1967 | 9.53333 | 0.66333333 | 0.631417726 | |
| | | | 33.55 | 23.53 | 23.55333333 | 9.99667 | 9.53333 | 0.46333333 | 0.725308503 | |
| | | Tobacco-5 | 27.19 | 25.53 | 25.56333333 | 1.62667 | 9.53333 | -7.9066667 | 239.9627511 | 251.2236388 |
| | | | 26.91 | 25.58 | 25.56333333 | 1.34667 | 9.53333 | -8.1866667 | 291.3615449 | |
| | | | 27.3 | 25.58 | 25.56333333 | 1.73667 | 9.53333 | -7.7966667 | 222.3466205 | |
| | | Tobacco-6 | 23.45 | 25.06 | 25.21666667 | -1.7667 | 9.53333 | -11.3 | 2521.383759 | 2607.737648 |
| | | | 23.24 | 25.13 | 25.21666667 | -1.9767 | 9.53333 | -11.51 | 2916.454801 | |
| | | | 23.53 | 25.46 | 25.21666667 | -1.6867 | 9.53333 | -11.22 | 2385.374385 | |
| 24.10.28 | MdGH3.1 | Sample | cp | Actin | Actin (Average | cp1 | cp2 | △cp | Relative gene expression | Relative gene expression (Average) |
| | | Tobacco (Control) | 33.64 | 20.51 | 20.73 | 12.91 | 13.5167 | -0.6066667 | 1.522736872 | 1.056627955 |
| | | | 34.29 | 20.72 | 20.73 | 13.56 | 13.5167 | 0.04333333 | 0.970410231 | |
| | | | 34.81 | 20.96 | 20.73 | 14.08 | 13.5167 | 0.56333333 | 0.676736762 | |
| | | Tobacco-7 | 26.72 | 21.26 | 21.62666667 | 5.09333 | 13.5167 | -8.4233333 | 343.3017338 | 195.7681223 |
| | | | 28.49 | 22.04 | 21.62666667 | 6.86333 | 13.5167 | -6.6533333 | 100.6590679 | |
| | | | 27.98 | 21.58 | 21.62666667 | 6.35333 | 13.5167 | -7.1633333 | 143.3435653 | |
| | | Tobacco-8 | 31.97 | 22.95 | 23.04333333 | 8.92667 | 13.5167 | -4.59 | 24.08394796 | 24.90035347 |
| | | | 31.67 | 23.1 | 23.04333333 | 8.62667 | 13.5167 | -4.89 | 29.65081798 | |
| | | | 32.17 | 23.08 | 23.04333333 | 9.12667 | 13.5167 | -4.39 | 20.96629446 | |
| | | Tobacco-9 | 34.39 | 20.8 | 20.83 | 13.56 | 13.5167 | 0.04333333 | 0.970410231 | 0.713398914 |
| | | | 35.01 | 20.89 | 20.83 | 14.18 | 13.5167 | 0.66333333 | 0.631417726 | |
| | | | 35.24 | 20.8 | 20.83 | 14.41 | 13.5167 | 0.89333333 | 0.538368784 | |
| | | Tobacco-10 | 33.7 | 26.35 | 26.6 | 7.1 | 13.5167 | -6.4166667 | 85.42975067 | 86.6796355 |
| | | | 34.65 | 26.97 | 26.6 | 8.05 | 13.5167 | -5.4666667 | 44.22121216 | |
| | | | 33.09 | 26.48 | 26.6 | 6.49 | 13.5167 | -7.0266667 | 130.3879437 | |
| | | Tobacco-11 | 35.73 | 24.93 | 24.93333333 | 10.7967 | 13.5167 | -2.72 | 6.588728138 | 11.75014373 |
| | | | 34.36 | 24.97 | 24.93333333 | 9.42667 | 13.5167 | -4.09 | 17.02992292 | |
| | | | 34.91 | 24.9 | 24.93333333 | 9.97667 | 13.5167 | -3.54 | 11.63178014 | |
| | | Tobacco-12 | 34.67 | 21.58 | 21.56666667 | 13.1033 | 13.5167 | -0.4133333 | 1.331759279 | 1.953131621 |
| | | | 33.94 | 21.5 | 21.56666667 | 12.3733 | 13.5167 | -1.1433333 | 2.208908001 | |
| | | | 33.87 | 21.62 | 21.56666667 | 12.3033 | 13.5167 | -1.2133333 | 2.318727582 | |
| | | Tobacco-13 | 35.65 | 23.62 | 23.61 | 12.04 | 13.5167 | -1.4766667 | 2.783049688 | 1.952139726 |
| | | | 36.14 | 23.48 | 23.61 | 12.53 | 13.5167 | -0.9866667 | 1.981601227 | |
| | | | 37 | 23.73 | 23.61 | 13.39 | 13.5167 | -0.1266667 | 1.091768265 | |
| | | Tobacco-14 | 34.25 | 25.2 | 25.31666667 | 8.93333 | 13.5167 | -4.5833333 | 23.97291323 | 20.89340023 |
| | | | 34.75 | 25.31 | 25.31666667 | 9.43333 | 13.5167 | -4.0833333 | 16.95140951 | |
| | | | 34.39 | 25.44 | 25.31666667 | 9.07333 | 13.5167 | -4.4433333 | 21.75587797 | |
| | | Tobacco-15 | 34.33 | 25.6 | 25.40666667 | 8.92333 | 13.5167 | -4.5933333 | 24.13965803 | 23.2955799 |
| | | | 34.56 | 25.31 | 25.40666667 | 9.15333 | 13.5167 | -4.3633333 | 20.58231471 | |
| | | | 34.27 | 25.31 | 25.40666667 | 8.86333 | 13.5167 | -4.6533333 | 25.16476697 | |
| 24.11.11 | Mdgh3.1 | Sample | cp | Actin | Actin (Average | cp1 | cp2 | △cp | Relative gene expression | Relative gene expression (Average) |
| | | Tobacco (Control) | 31.79 | 20.19 | 20.33666667 | 11.4533 | 11.1167 | 0.33666667 | 0.791868805 | 1.03723411 |
| | | | 31.65 | 20.35 | 20.33666667 | 11.3133 | 11.1167 | 0.19666667 | 0.872564288 | |
| | | | 30.92 | 20.47 | 20.33666667 | 10.5833 | 11.1167 | -0.5333333 | 1.447269237 | |
| | | Tobacco-16 | 31.07 | 21.58 | 21.59 | 9.48 | 11.1167 | -1.6366667 | 3.109465621 | 3.271871128 |
| | | | 31.07 | 21.64 | 21.59 | 9.48 | 11.1167 | -1.6366667 | 3.109465621 | |
| | | | 30.86 | 21.55 | 21.59 | 9.27 | 11.1167 | -1.8466667 | 3.596682143 | |
| | | Tobacco-17 | 30.15 | 21.16 | 21.20333333 | 8.94667 | 11.1167 | -2.17 | 4.500233939 | 3.698220311 |
| | | | 30.54 | 21.34 | 21.20333333 | 9.33667 | 11.1167 | -1.78 | 3.434261746 | |
| | | | 30.66 | 21.11 | 21.20333333 | 9.45667 | 11.1167 | -1.66 | 3.160165247 | |
| | | Tobacco-18 | 31.33 | 21.8 | 21.84333333 | 9.48667 | 11.1167 | -1.63 | 3.095129987 | 3.793198171 |
| | | | 30.79 | 21.69 | 21.84333333 | 8.94667 | 11.1167 | -2.17 | 4.500233939 | |
| | | | 31.04 | 22.04 | 21.84333333 | 9.19667 | 11.1167 | -1.92 | 3.784230587 | |
| | | Tobacco-19 | 31.84 | 20.9 | 20.96 | 10.88 | 11.1167 | -0.2366667 | 1.178267139 | 1.324952756 |
| | | | 31.32 | 20.97 | 20.96 | 10.36 | 11.1167 | -0.7566667 | 1.689582347 | |
| | | | 31.93 | 21.01 | 20.96 | 10.97 | 11.1167 | -0.1466667 | 1.107008782 | |
| | | Tobacco-20 | 31.89 | 25.65 | 25.56 | 6.33 | 11.1167 | -4.7866667 | 27.60134502 | 42.48854302 |
| | | | 31.26 | 25.5 | 25.56 | 5.7 | 11.1167 | -5.4166667 | 42.71487533 | |
| | | | 30.84 | 25.53 | 25.56 | 5.28 | 11.1167 | -5.8366667 | 57.14940871 | |
| | | Tobacco-21 | 31.07 | 21.01 | 21.14333333 | 9.92667 | 11.1167 | -1.19 | 2.281527432 | 2.434555628 |
| | | | 30.45 | 21.26 | 21.14333333 | 9.30667 | 11.1167 | -1.81 | 3.506422885 | |
| | | | 31.66 | 21.16 | 21.14333333 | 10.5167 | 11.1167 | -0.6 | 1.515716567 | |

**Supplementary table S2.** Quantitative analysis of the relative expression levels of MdGH3.1 in transgenic tobacco. The control is wild-type tobacco. The color of the cells indicates the extent of MdGH3.1 overexpression. Blue cells indicate relative expression levels between 2 and 10, considered low overexpression. Green cells indicate relative expression levels greater than 20, considered high overexpression. Yellow cells indicate MdGH3.1 overexpression.

| Time | Gene | Sample | cp | Actin | Actin (Average) | cp1 | cp2 | △cp | Relative gene expression | Relative gene expression (Average) |
|---|---|---|---|---|---|---|---|---|---|---|
| | | | 30.78 | 23.8 | 23.89333333 | 6.886666667 | 6.636666667 | 0.25 | 0.840896415 | |
| | | Arabidopsis （Control） | 30.59 | 23.96 | 23.89333333 | 6.696666667 | 6.636666667 | 0.06 | 0.959264119 | 1.013289412 |
| | | | 30.22 | 23.92 | 23.89333333 | 6.326666667 | 6.636666667 | -0.31 | 1.2397077 | |
| | | | 27.81 | 25.63 | 25.75 | 2.06 | 6.636666667 | -4.576666667 | 23.86239041 | |
| | MdGH3.1 | Arabidopsis -1 | 27.7 | 25.9 | 25.75 | 1.95 | 6.636666667 | -4.686666667 | 25.75296552 | 20.38004361 |
| | | | 28.86 | 25.72 | 25.75 | 3.11 | 6.636666667 | -3.526666667 | 11.52477489 | |
| | | | 31.71 | 26.06 | 26.16333333 | 5.546666667 | 6.636666667 | -1.09 | 2.128740365 | |
| | | Arabidopsis -2 | 31.8 | 26.34 | 26.16333333 | 5.636666667 | 6.636666667 | -1 | 2 | 3.341939078 |
| | | | 30.24 | 26.09 | 26.16333333 | 4.076666667 | 6.636666667 | -2.56 | 5.897076869 | |
| | | | 26.03 | 20.81 | 20.82333333 | 5.206666667 | 5.276666667 | -0.07 | 1.049716684 | |
| | | Arabidopsis （Control） | 26.22 | 20.89 | 20.82333333 | 5.396666667 | 5.276666667 | 0.12 | 0.920187651 | 1.001723086 |
| | | | 26.05 | 20.77 | 20.82333333 | 5.226666667 | 5.276666667 | -0.05 | 1.035264924 | |
| | | | 29.45 | 27.59 | 27.74333333 | 1.706666667 | 5.276666667 | -3.57 | 11.87618857 | |
| 24.10.10 | MdGH3.1 | Arabidopsis -3 | 30.12 | 27.77 | 27.74333333 | 2.376666667 | 5.276666667 | -2.9 | 7.464263932 | 9.66230504 |
| | | | 29.75 | 27.87 | 27.74333333 | 2.006666667 | 5.276666667 | -3.27 | 9.646462622 | |
| | | | 30.43 | 26.02 | 26.12 | 4.31 | 5.276666667 | -0.966666667 | 1.954319937 | |
| | | Arabidopsis -4 | 29.56 | 26.2 | 26.12 | 3.44 | 5.276666667 | -1.836666667 | 3.571838044 | 2.976275542 |
| | | | 29.63 | 26.14 | 26.12 | 3.51 | 5.276666667 | -1.766666667 | 3.402668644 | |
| | | | 29.51 | 25.05 | 25.20333333 | 4.306666667 | 3.843333333 | 0.463333333 | 0.725308503 | |
| | | Arabidopsis （Control） | 28.65 | 25.27 | 25.20333333 | 3.446666667 | 3.843333333 | -0.396666667 | 1.316462719 | 1.029688448 |
| | | | 28.98 | 25.29 | 25.20333333 | 3.776666667 | 3.843333333 | -0.066666667 | 1.047294123 | |
| | | | 29.2 | 26.18 | 26.20666667 | 2.993333333 | 3.843333333 | -0.85 | 1.802500925 | |
| | MdGH3.1 | Arabidopsis -5 | 29.49 | 26.21 | 26.20666667 | 3.283333333 | 3.843333333 | -0.56 | 1.474269217 | 1.947873979 |
| | | | 28.69 | 26.23 | 26.20666667 | 2.483333333 | 3.843333333 | -1.36 | 2.566851795 | |
| | | | 28.79 | 24.74 | 24.75333333 | 4.036666667 | 3.843333333 | 0.193333333 | 0.87458267 | |
| | | Arabidopsis -6 | 28.78 | 24.67 | 24.75333333 | 4.026666667 | 3.843333333 | 0.183333333 | 0.880665874 | 0.840637572 |
| | | | 28.98 | 24.85 | 24.75333333 | 4.226666667 | 3.843333333 | 0.383333333 | 0.766664172 | |
| Time | Gene | Sample | cp | Actin | Actin (Average) | cp1 | cp2 | △cp | Relative gene expression | Relative gene expression (Average) |
| | | | 33.78 | 28.78 | 29.00666667 | 4.773333333 | 4.526666667 | 0.246666667 | 0.842841545 | |
| | | Arabidopsis （Control） | 33.41 | 29.06 | 29.00666667 | 4.403333333 | 4.526666667 | -0.123333333 | 1.089248656 | 1.007112952 |
| | | | 33.41 | 29.18 | 29.00666667 | 4.403333333 | 4.526666667 | -0.123333333 | 1.089248656 | |
| | | | 33.46 | 32.8 | 32.74 | 0.72 | 4.526666667 | -3.806666667 | 13.99332272 | |
| | | Arabidopsis -7 | 33.25 | 31.77 | 32.74 | 0.51 | 4.526666667 | -4.016666667 | 16.18591104 | 14.7241855 |
| | | | 33.46 | 33.65 | 32.74 | 0.72 | 4.526666667 | -3.806666667 | 13.99332272 | |
| | | | 32.36 | 30.87 | 31.10333333 | 1.256666667 | 4.526666667 | -3.27 | 9.646462622 | |
| 24.10.16 | MdGH3.1 | Arabidopsis -8 | 33.47 | 31.17 | 31.10333333 | 2.366666667 | 4.526666667 | -2.16 | 4.469148552 | 6.174410546 |
| | | | 33.49 | 31.27 | 31.10333333 | 2.386666667 | 4.526666667 | -2.14 | 4.407620464 | |
| | | | 32.89 | 23.12 | 23.27 | 9.62 | 4.526666667 | 5.093333333 | 0.029292328 | |
| | | Arabidopsis -9 | 34.19 | 23.2 | 23.27 | 10.92 | 4.526666667 | 6.393333333 | 0.011896382 | 0.015554054 |
| | | | 35.31 | 23.49 | 23.27 | 12.04 | 4.526666667 | 7.513333333 | 0.005473452 | |
| | | | 35.2 | 26.55 | 26.76666667 | 8.433333333 | 4.526666667 | 3.906666667 | 0.066677015 | |
| | | Arabidopsis -10 | 35.72 | 26.89 | 26.76666667 | 8.953333333 | 4.526666667 | 4.426666667 | 0.046498672 | 0.134338878 |
| | | | 33.08 | 26.86 | 26.76666667 | 6.313333333 | 4.526666667 | 1.786666667 | 0.289840948 | |
| Time | Gene | Sample | cp | Actin | Actin (Average) | cp1 | cp2 | △cp | Relative gene expression | Relative gene expression (Average) |
| | | | 34.36 | 19.94 | 20.04666667 | 14.31333333 | 13.14666667 | 1.166666667 | 0.445449359 | |
| | | Arabidopsis （Control） | 31.74 | 20.1 | 20.04666667 | 11.69333333 | 13.14666667 | -1.453333333 | 2.738400258 | 1.334547872 |
| | | | 33.48 | 20.1 | 20.04666667 | 13.43333333 | 13.14666667 | 0.286666667 | 0.819793998 | |
| | | | 32.48 | 23.71 | 23.81333333 | 8.666666667 | 13.14666667 | -4.48 | 22.31589866 | |
| | | Arabidopsis -11 | 32.83 | 23.53 | 23.81333333 | 9.016666667 | 13.14666667 | -4.13 | 17.50869922 | 15.7286997 |
| | | | 34.08 | 24.2 | 23.81333333 | 10.26666667 | 13.14666667 | -2.88 | 7.361501205 | |
| | | | 33.74 | 23.06 | 23.03666667 | 10.70333333 | 13.14666667 | -2.443333333 | 5.438969491 | |
| 24.10.18 | MdGH3.1 | Arabidopsis -12 | 33.19 | 22.99 | 23.03666667 | 10.15333333 | 13.14666667 | -2.993333333 | 7.963117433 | 5.671809242 |
| | | | 34.33 | 23.06 | 23.03666667 | 11.29333333 | 13.14666667 | -1.853333333 | 3.613340803 | |
| | | | 31.71 | 23.01 | 22.93 | 8.78 | 13.14666667 | -4.366666667 | 20.62992494 | |
| | | Arabidopsis -13 | 34.74 | 22.9 | 22.93 | 11.81 | 13.14666667 | -1.336666667 | 2.525670902 | 9.449650063 |
| | | | 33.7 | 22.88 | 22.93 | 10.77 | 13.14666667 | -2.376666667 | 5.193354353 | |
| | | | 32.12 | 23.63 | 23.75333333 | 8.366666667 | 13.14666667 | -4.78 | 27.47409397 | |
| | | Arabidopsis -14 | 32.93 | 23.83 | 23.75333333 | 9.176666667 | 13.14666667 | -3.97 | 15.67072476 | 21.22654436 |
| | | | 32.54 | 23.8 | 23.75333333 | 8.786666667 | 13.14666667 | -4.36 | 20.53481436 | |

**Supplementary table S3.** Quantitative analysis of the relative expression levels of MdGH3.1 in transgenic *Arabidopsis thaliana*. The control is wild-type *Arabidopsis thaliana*. Yellow cells indicate MdGH3.1 overexpression.
